# Manganese suppresses tumor growth through hyper-activating IRE1α

**DOI:** 10.1101/2025.03.16.643497

**Authors:** Ruoxi Shi, Ming Wang, Si Chen, Xinwei Hu, Xudong Zhao, Shixin Zhou, Congli Fan, Zihan Zhou, Likun Wang

**Affiliations:** National Laboratory of Biomacromolecules, CAS Center for Excellence in Biomacromolecules, Institute of Biophysics, Chinese Academy of Sciences, 100101 Beijing, P.R. China; College of Life Sciences, University of Chinese Academy of Sciences, 100049 Beijing, P.R. China

**Author notes:** These authors contributed equally (Ruoxi Shi, Ming Wang).

## Abstract

IRE1α and its downstream XBP1 signal is the most conserved unfolded protein response pathway that cells utilize to combat endoplasmic reticulum stress, also known to be utilized by tumor cells to adapt to harsh environment, leading to tumor progression. Several inhibitors against IRE1α have been developed, some of which show promising effect in clinical trial for cancer therapy, but none of them have been used in practice. Considering that hyper-activation of IRE1α induces cell death, we hypothesize that activation of IRE1α could be an alternative way for tumor suppression. Here, we identified divalent manganese ion as a potent activator to IRE1α, which interacts with the cytosolic part of IRE1α directly, augmenting the downstream pro-apoptotic pathway but not the pro-survival outcome. Mn^2+^ limits tumor growth in xenograft model in an IRE1α-dependent way. Our finding suggests pharmacological activation of IRE1α as an underestimated but promising way in cancer therapy.

**Highlights:**

- Mn^2+^ binds to cytosolic segment of IRE1α and promotes its oligomerization
- Mn^2+^ hyper-activates IRE1α and induces downstream terminal UPR under ER stress
- Mn^2+^ does not elevate XBP1s expression
- Mn^2+^ as an IRE1α activator impairs tumor growth in xenograft model

## Introduction

Endoplasmic reticulum (ER) stress is a condition characterized by the accumulation of misfolded and unfolded proteins in the ER lumen that exceeds the protein processing capacity. Under ER stress, cells initiate the unfolded protein response (UPR) to restore ER homeostasis^1^. Genetic mutation, metabolic abnormality, chemotherapy, among others, constitutively challenge the protein folding in tumor cells to cause ER stress^2^. Activation of the tumoral UPR is notorious for its contribution to tumor progression by building up cell adaptation, bypassing antitumor immunity, and facilitating angiogenesis and metastasis^3^.

As the most conserved UPR pathway, IRE1α has been extensively studied for its role in tumor progression^4^. IRE1α is an ER-transmembrane kinase and RNase which, upon ER stress, undergoes dimerization/oligomerization and trans-autophosphorylation, activating the RNase domain to initiate the mRNA splicing of XBP1^5^. Removal of a 26-nucleotide intron results in the translation of XBP1s (s, spliced), a transcription factor that plays key roles in cell adaptation^6–8^. In many cancers, IRE1α-XBP1 signal positively correlates with tumor malignancy and presages poor survival of the patients. Thus, IRE1α-XBP1 has been regarded as a potential target for cancer therapy. Many IRE1α inhibitors have been developed^4^. However, none of them have been used clinically.

Although IRE1α-XBP1 serves to maintain ER homeostasis, unmitigated IRE1α activation can also elicit cell death, especially during severe and chronic ER stress. This has been studied extensively, ranging from IRE1α’s role in mRNA and microRNA degradation (termed regulated IRE1α-dependent decay) to activation of JNK phosphorylation^9–11^. Considering that tumor tissues experience ER stress and have basal level of IRE1α activation, we proposed that forced hyper-activation of IRE1α could be an alternative way in cancer therapy. However, very few has been done in developing IRE1α activators. A recent study reported a high-throughput screening that identified IXA4 as a small compound to activate IRE1-XBP1 pathway, yet the molecular target of the compound is still unclear ^12^. CRUK-3 and quercetin can also activate IRE1α RNase activity, but both act as kinase inhibitors ^13,14^.

Here, we reported that divalent Mn ion (Mn^2+^) causes cell death in the context of ER stress in an IRE1α-dependent manner. by upregulating the activity of IRE1α and the downstream pro-apoptotic signals. Mn^2+^ promotes IRE1α phosphorylation and oligomerization, and activates the downstream pro-apoptotic signals yet with the protective XBP1 pathway unaffected. Mn^2+^ administration repressed tumor progression in mice experiment in an IRE1α-dependent manner, suggesting that Mn^2+^-driven hyperactivation of IRE1α could be a potential strategy for cancer therapy.

## Results

### Mn^2+^ impairs cell viability and increases ER stress-induced cell death in an IRE1α-dependent manner

Divalent ions play essential roles in cellular function and are tightly linked with the pathology of many diseases. In search of divalent ions that may hyper-activate IRE1α to a level that hinder tumor cell growth, we found that Mn^2+^ impaired cell viability and aggravated cell death in MDA-MB-231, a human triple-negative breast cancer (TNBC) cell line. We chose MDA-MB-231 because it has intrinsically highly activated IRE1α, which we think may be more prone to be hyper-activated^15^. Mn^2+^ had no effect on cell viability nor caused cell death by itself, but showed cytotoxicity in a dose-dependent manner in the presence of thapsigargin (Tg), an ER stress inducer (**Figures 1A and 1B**). Notably, the cytotoxic effect is ion-type specific, since other essential divalent ions, including Ca^2+^, Mg^2+^, Zn^2+^, and Cu^2+^, did not decrease cell viability in the context of ER stress (**Figures S1A and S1B**). In addition, induction of ER stress by culturing cells under oxygen-glucose deprivation (OGD) also rendered cells to be sensitive to Mn^2+^ insult (**Figure S1C**).

**Figure 1.**
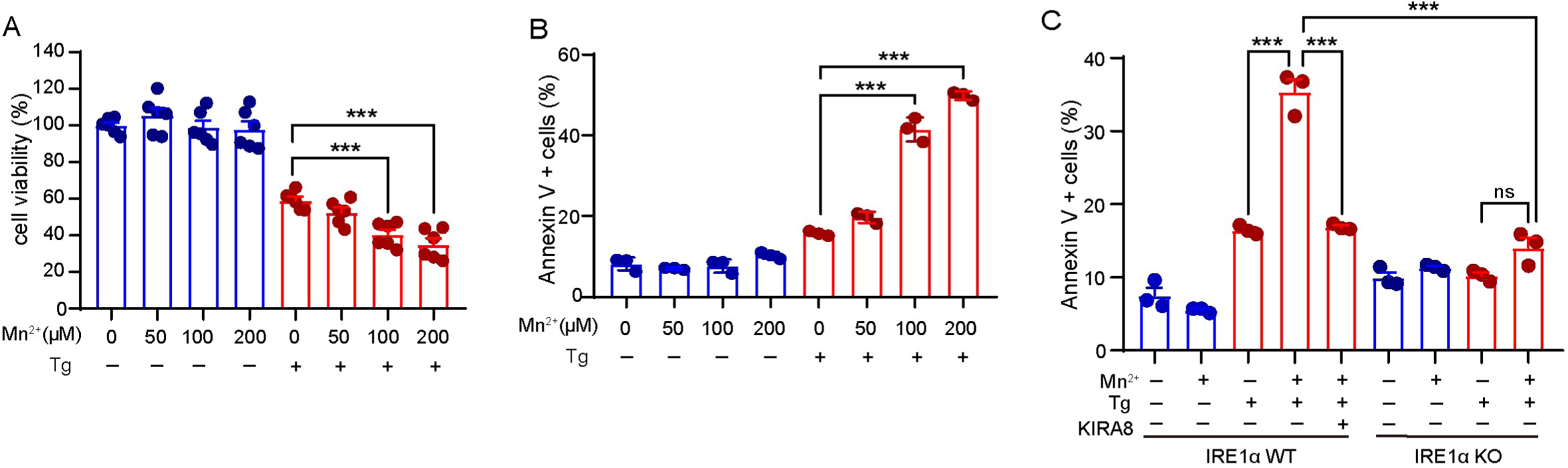
Mn^2+^ impairs cell viability and increases ER stress-induced cell death in an IRE1α-dependent manner. (A) Cell viability detected by the MTT assay. MDA-MB-231 cells were treated with or without Tg (0.2 μM) in the absence or presence of different concentrations of MnCl_2_ for 24 h (n = 6). (**B**) Cell death percentage in MDA-MB-231 cells measured by Annexin V staining. Cells were treated as in (**A**). The percentage of Annexin V-positive cells was shown (n = 3). (**C**) Cell death percentage in MDA-MB-231 WT and IRE1α KO cells measured by Annexin V staining. Cells were treated with Tg (0.2 μM), MnCl_2_ (100 μM) and KIRA8 (1 μM) alone or in combination for 24 h (n = 3). Data are presented as mean ± SEM. Statistical analysis was performed by unpaired Student’s t test. ns, not significant; ***p < 0.001.

The cytotoxic effect of Mn^2+^ was only observed in the presence of ER stress, suggesting that a basal activation of the UPR is a prerequisite. We hypothesized that Mn^2+^ enhanced the activity of the UPR signals to exceed a threshold, exerting the pro-apoptotic function. Constitutive overexpression of IRE1α has been reported to induce cell apoptosis^11^. Interestingly, pharmacological inhibition of IRE1α pathway with KIRA8^16,17^, or genetic deletion of IRE1α, reduced the cell apoptotic level, suggesting that IRE1α activation could be a reason for cell death (**Figures 1C and S1C**).

### Mn^2+^ promotes the terminal UPR downstream of IRE1α but spares the adaptive UPR unaffected

To elucidate whether Mn^2+^ activates IRE1α, we then examined IRE1α pathway activity in ER-stressed cells in the presence or absence of Mn^2+^. Upon activation, IRE1α undergoes dimerization and oligomerization, and becomes auto-phosphorylated. We found that Mn^2+^ led to a shift of endogenous IRE1α to the high-molecular-weight fractions (**Fig. 2A**). To visualize the oligomerization of IRE1α, we expressed a GFP-tagged IRE1α in the cell, and examined the puncta formation upon Tg treatment. Tg induced puncta formation as expected. Importantly, this was further boosted by Mn^2+^ (**Fig. 2B**). In line with this, Mn^2+^ advanced the Tg-induced IRE1α phosphorylation and the downstream *XBP1* splicing (**Figures 2C-F**). This was also the case for cells under OGD condition, where Mn^2+^ promotes both IRE1α phosphorylation and XBP1 splicing (**Figures S2A-D**). Notably, Mn^2+^ itself did not promote IRE1α oligomerization, and had no effect on IRE1α phosphorylation or *XBP1* splicing (**Figures 2B-F and S2A-D**). On the other hand, Mn^2+^ was able to increase the phosphorylation level of over-expressed IRE1α that was intrinsically phosphorylated, even in the absence of ER stress (**Figures 2G and 2H**). Thus, Mn^2+^ serves as an amplifier to increase the activity of IRE1α once IRE1α is partially activated.

**Figure 2.**
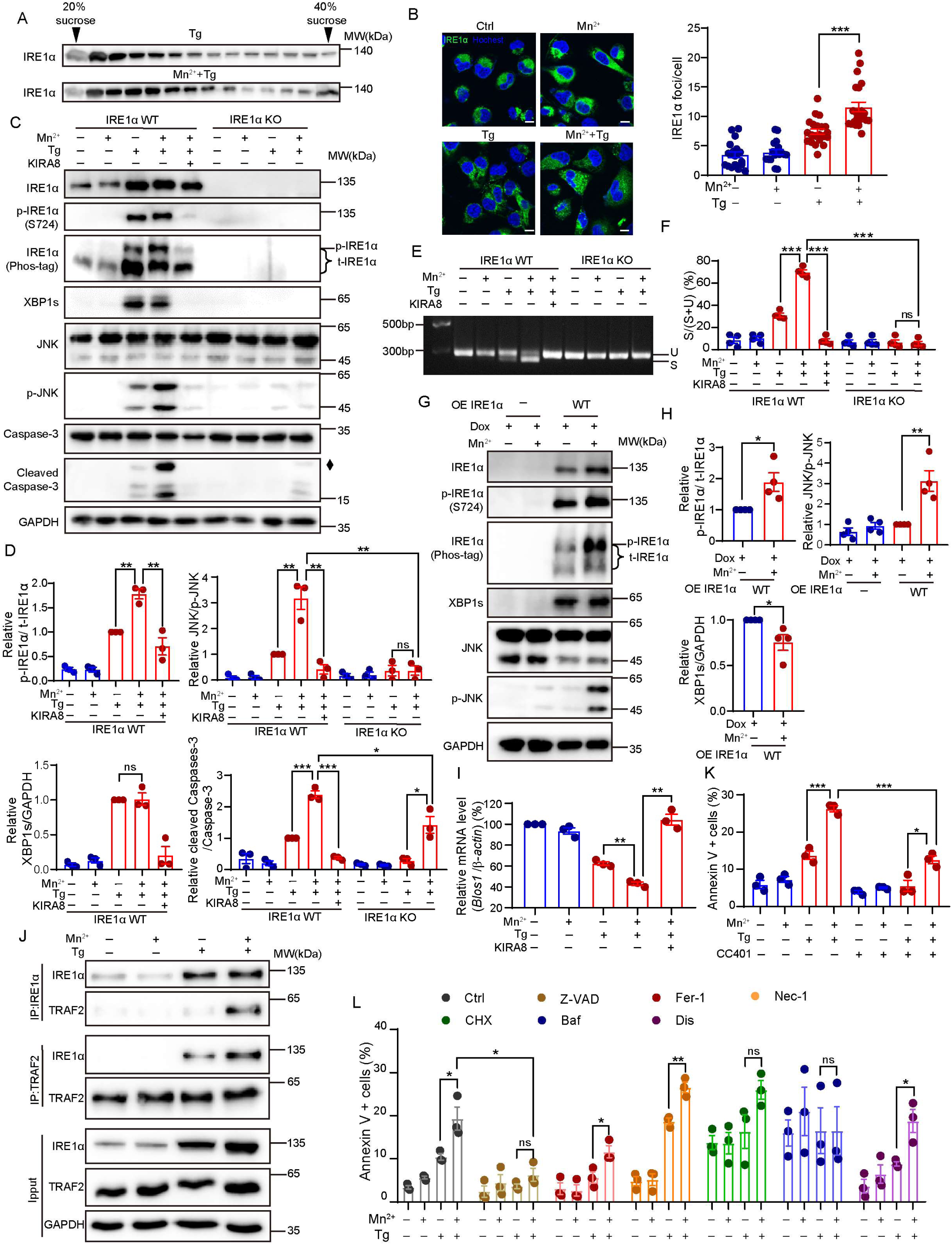
Mn^2+^ promotes the terminal UPR downstream of IRE1α but spares the adaptive UPR unaffected. (**A**) The oligomerization of IRE1α detected by immunoblotting assay. MDA-MB-231 cells were treated with Tg (0.2 μM) and MnCl_2_ (100 μM) alone or in combination for 24 h. Cell lysates were separated by sucrose density gradient centrifugation (20%–40%). (**B**) Representative immunofluorescence image (Left) and the quantitative result (Right) of hIRE1α-sfGFP puncta in MDA -MB-231 IRE1α KO cells that stably express hIRE1α-sfGFP cells in a Dox-inducible way. Cells were pretreated with Dox (5 μg/mL) for 20 h, followed by MnCl_2_ (100 μM) treatment for 4 h. Hoechst is used to identify the nucleus. Scale bar: 10 μm. (**C**) Phosphorylation of IRE1α and activation of its downstream signaling pathway in MDA-MB-231 WT and IRE1α KO cells measured by immunoblotting assay. Cells were treated with Tg (0.2 μM), MnCl_2_ (100 μM) and KIRA8(1 μM) alone or in combination for 24 h. Diamond denotes non-specific band. (**D**) Quantitative gray value analysis of p-IRE1α/t-IRE1α (by Phos-tag), p-JNK/t-JNK, XBP1s/GAPDH, and cleaved Caspase-3/Caspase-3 in (**C**) (n = 3). Relative ratio was shown, with ratio in lane 3 (Tg treatment for IRE1α WT) set as 1. (**E**) Agarose gel of *Xbp1* cDNA amplicons from MDA-MB-231 WT and IRE1α KO cells. Cells were treated as in (**C**). U, unspliced *Xbp1*; S, spliced *Xbp1*. (**F**) Quantitative gray value analysis of S/(S+U) in (**E**) (n = 4). (**G**) Immunoblotting assay of IRE1α phosphorylation and its downstream signaling pathways in MDA-MB-231 IRE1α KO cells that stably express exogenous IRE1α in a Dox-inducible way. Cells were pretreated with Dox (200 ng/mL) for 24 h, and then incubated with MnCl_2_ (200 μM) for 48 h. Relative ratio was shown, with ratio in lane 3 (OE IRE1α WT) set as 1. (**H**) Quantitative gray value analysis of p-IRE1α/t-IRE1α (by Phos-tag), p-JNK/t-JNK, and XBP1s/GAPDH in (**G**) (n = 4). (**I**) The mRNA expression level of *Blos1* in MDA-MB-231 cells detected by RT-qPCR. Cells were pretreated with ActD (2 μg/mL) for 1 h, then treated with Tg (0.2 μM), MnCl_2_ (100 μM) and KIRA8 (1μM) alone or in combination for 24 h (n = 3). (**J**) The interaction of IRE1α and TRAF2 in MDA-MB-231 cells detected by co-immunoprecipitation. Cells were treated with Tg (0.2 μM) and MnCl_2_ (100 μM) alone or in combination for 24 h. (**K**) Cell death percentage in MDA-MB-231 cells measured by Annexin V staining. Cells were pretreated with or without CC401 (5 μM) for 1 h, then incubated with Tg (0.2 μM) and MnCl_2_ (100 μM) alone or in combination for 24 h (n = 3). (**L**) Cell death percentage in MDA-MB-231 cells measured by Annexin V staining. Cells were pretreated with z-VAD-FMK (z-VAD, 20 μM), ferrostatin-1 (Fer, 1 μM), necrostatin-1 (Nec, 80 μM), bafilomycin A1 (Baf, 10 nM), disulfiram (Dis, 10 μM) for 24 h or cycloheximide (CHX, 2 μM) for 6 h, and further treated with Tg (0.2 μM) and MnCl_2_ (100 μM) alone or in combination for 24 h (n = 3). Data are presented as mean ± SEM. Statistical analysis was performed by unpaired Student’s t test. ns, not significant; *p < 0.05; **p < 0.01; ***p < 0.001.

In cells that experience ER stress, IRE1α plays dual roles in cell fate decision. On one hand, IRE1α serves as an RNase to initiate xbp1 mRNA splicing, resulting in the translation of a transcription factor XBP1s that restores the ER homeostasis. On the other hand, persistent activation of IRE1α decreases the RNA substrate selectivity and results in rampant degradation of certain mRNAs and micro-RNAs, leading to cell dysfunction and programmed cell death^9,10,18^. IRE1α could also interact with TRAF2, activating the downstream pro-apoptotic JNK signaling^19^. We found that Mn^2+^ failed to elevate the protein amount of XBP1s despite an increased *xbp1* splicing ratio in both Tg- and OGD-treated cells (**Figures 2C-F and S2A-D**). Similarly, in IRE1α-overexpressing cells that had XBP1s expression induced, Mn^2+^ was unable to further increase XBP1s protein level (**Figures 2G and 2H**). Considering the bona fide effect of XBP1s, this result implied that Mn^2+^-induced IRE1α activation may not benefit cells in combatting ER stress.

Rather, Mn^2+^ aggravated the Tg- or OGD-induced mRNA level decrease for *Blos1*, a known RNA substrate of IRE1α, indicating an increased level of RIDD^20^. The decrease in *Blos1* could be suppressed by IRE1α inhibition (**Figure 2I and S2E**). Furthermore, Mn^2+^ significantly promoted IRE1α-TRAF2 interaction and advanced the JNK phosphorylation and the downstream caspase-3 cleavage in Tg-treated cells but not naïve cells (**Figures 2C, 2D, and 2J**). Both JNK phosphorylation and caspase-3 cleavage were suppressed upon IRE1α inhibition or deletion, supporting the idea that Mn^2+^ activated JNK signaling through IRE1α (**Figure 2C and 2D**). In addition, inhibition of JNK with CC401 significantly reduced the apoptotic level in Mn^2+^- and Tg-treated MDA-MB-231 cells, consistent with its pro-apoptotic role (**Figure 2K**). We further examined the contribution of different type of cell death to the cytotoxicity of Mn^2+^ by inhibiting various types of cell death. Z-VAD, pan caspase inhibitor, significantly reduced cell death that was triggered by Tg and aggravated by Mn^2+^ (**Figure 2L**). In cells experiencing OGD, Mn^2+^ also promoted JNK phosphorylation and caspase-3 cleavage in an IRE1α-dependent manner (**Supplementary Figures S2A and S2B**). Similarly, Mn^2+^ increased cell death under OGD treatment, which can be alleviated by JNK inhibition or caspase inhibition (**Supplementary Figures S2F and S2G**). Finally, while IRE1α overexpressing per se was unable to induce JNK phosphorylation, this pro-death signal became pronounced upon Mn^2+^ treatment (**Figures 2G and 2H**). Thus, Mn^2+^ promotes JNK- and caspase-dependent cell death whenever IRE1α is partially activated by further activating IRE1α and its downstream terminal UPR but not adaptive UPR.

### Mn^2+^ interacts with the cytosolic domain of IRE1_α_, and promotes the oligomerization and auto-phosphorylation of IRE1α

We wondered whether Mn^2+^ directly activated IRE1α. To this end, we expressed both the luminal domain (IRE1α(LD), residues 24-441) and cytosolic fractions of IRE1α (IRE1α(CD), residues 469-977), and measured the protein melting curve in the presence or absence of Mn^2+^ by thermal shift assay (TSA) (**Figures S3A and S3B**). Mn^2+^ increased the melting temperature of IRE1α(CD), but not IRE1α(LD), in a concentration-dependent manner, indicating a direct interaction between Mn^2+^ and the cytosolic domain of IRE1α (**Figures 3A, 3B, S3C, and S3D**). Furthermore, Mn^2+^ increased the phosphorylation level of IRE1α(CD), which was even not observed with same concentration of Mg^2+^, suggesting that Mn^2+^ may be a more potent cofactor than Mg^2+^ for IRE1α kinase activity (**Figure 3C and 3D**). Not only this, Mn^2+^ augmented IRE1α(CD) dimerization, increased the IRE1α(CD) RNase activity towards a synthesized *xbp1* RNA fragment that harbors the cleavage site for IRE1α, and promoted the in vitro degradation of mouse Ins2 mRNA by IRE1α(CD) (**Figures 3E-3H**). These in vitro experiments demonstrated that Mn^2+^ directly binds to and activates IRE1α.

**Figure 3.**
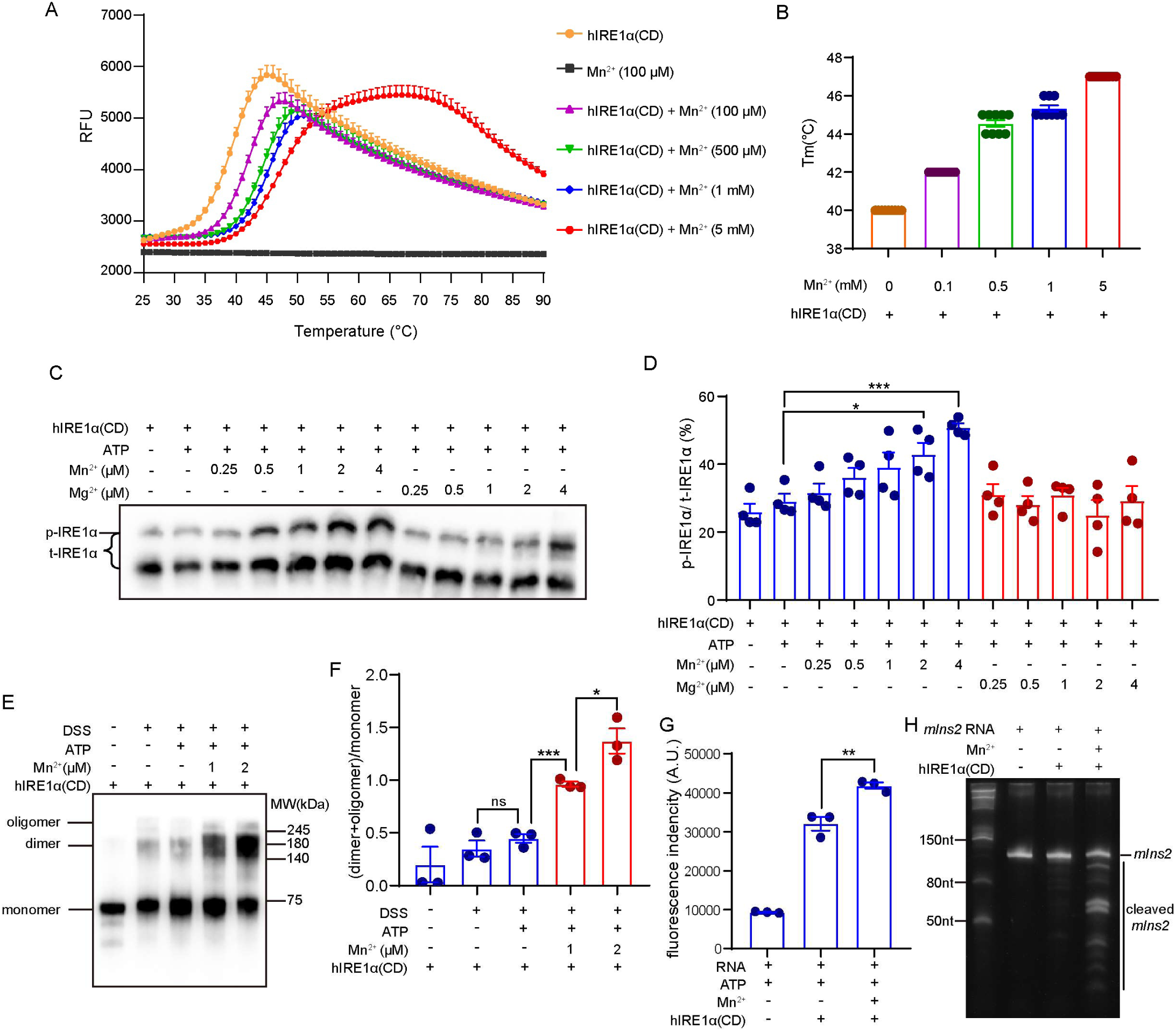
Mn^2+^ interacts with the cytosolic domain of IRE1α, and promotes the oligomerization and auto-phosphorylation of IRE1α. (**A**) The unfolding curve of recombinant hIRE1α(CD) (residues 469-977) protein (2 μM) incubated with different concentrations of MnCl_2_ measured by thermal shift assay. The melting temperature (Tm) corresponds to the horizontal axis value at the point of maximum slope (n = 9). (**B**) Comparison of the melting temperature of hIRE1α(CD) incubated with MnCl_2_ (n = 9). (**C**) The phosphorylation of hIRE1α(CD) was detected by Phos-tag immunoblotting assay. hIRE1α(CD) protein was incubated with different concentrations of Mn^2+^ or Mg^2+^ at room temperature for 2 h. (**D**) Quantitative analysis of p-IRE1α/t-IRE1α (by Phos-tag) in (**C**) (n = 4). (**E**) The oligomerization of hIRE1α(CD) was detected by immunoblotting assay. hIRE1α(CD) (10 μM) was incubated with or without MnCl_2_ (1 or 2 mM) for 90 min at room temperature, then crosslinked by adding 250 μM disuccinimidyl suberate for 30 min. (**F**) Quantitative analysis of (dimer+oligomer)/monomer in (**E**) (n = 3). (**G**) The RNase activity of hIRE1α(CD) was detected by the in vitro cleavage assay. hIRE1α(CD) (1 μM) protein was incubated with MnCl_2_ (1 mM) for 20 min at room temperature, followed by incubation with 3 μM RNA substrate for 5 min. The reaction was quenched by adding urea to a final concentration of 4 M. The fluorescence of mini-XBP1 was detected using a microplate reader (n = 3). (**H**) The degradation of mouse *Ins2* mRNA (*mIns2*) was detected by the TBE urea polyacrylamide gel. hIRE1α(CD) (5 μM) was incubated with in vitro-transcribed RNA of *mIns2* (110 nt) (0.5 μM) and MnCl_2_ (2 mM) at 30 °C for 2 h. Data are presented as mean ± SEM. Statistical analysis was performed by unpaired Student’s t test. ns, not significant; *p < 0.05; **p < 0.01; ***p < 0.001.

### Mn^2+^ impairs tumor growth in xenograft model

Next, we examined the effect of Mn^2+^ on tumor growth in mice with MDA-MB-231 (either WT or IRE1α-KO) xenotransplantation. Injection of Mn^2+^ impaired WT tumor growth, but could not limit IRE1α-KO tumor growth (**Figures 4A-4D**). Mn^2+^ had no effect on mice weight (**Figure 4E**). We found that Mn^2+^ administration increased the phosphorylation level of IRE1α and JNK (**Figures 4F and 4G**). Interestingly, XBP1 protein level was not increased, but rather decreased upon Mn^2+^ treatment, indicating again that Mn^2+^ did not promote the pro-survival signal downstream of IRE1α (**Figures 4F and 4G**). As for IRE1α-KO tumors, we did not see upregulation of JNK phosphorylation after Mn^2+^ treatment, which was instead decreased (**Figure 4H**). Although the reason remains unknown, such phenomenon is consistent with the observation that Mn^2+^ slightly promoted IRE1α-KO tumor growth (**Figures 4B-4D**). Finally, cell apoptosis in tumor tissue was examined by TUNEL staining. As expected, Mn^2+^ treatment increased cell apoptosis in WT, but not IRE1α-KO tumors, demonstrating that Mn^2+^ promotes tumor cell death and retards tumor growth through activating IRE1α (**Figures 4I and 4J**).

**Figure 4.**
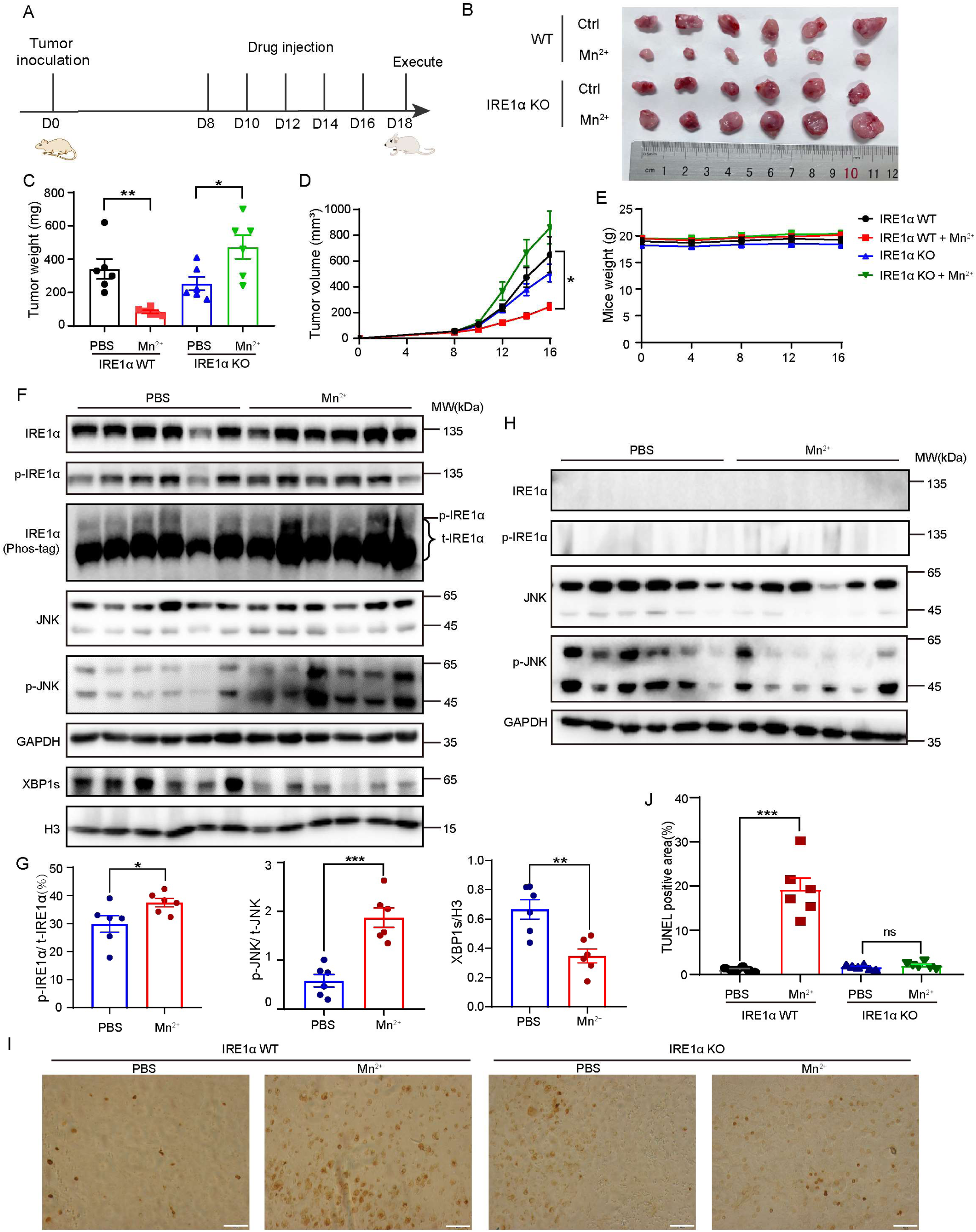
Mn^2+^ impairs tumor growth in xenograft model. (**A**) The schematic of the MnCl_2_ treatment schedule. BALB/c nude mice were subcutaneously inoculated with MDA-MB-231 WT or IRE1α KO cells (1.5×10^6^ cells per mouse), and peritumorally injected with saline or MnCl_2_ (5 mg/kg) every two days from day 8. Tumors were collected on day 18 after mice execution. (**B**) - (**E**) Photograph of tumor (**B**) and tumor weight (**C**) measured after mice execution. Tumor volume (**D**) and the body weight (**E**) of mice inoculated with MDA-MB-231 WT or IRE1α KO cells during drug injection were recorded (n = 6). (**F**) Immunoblotting assay of IRE1α phosphorylation and its downstream signaling pathways in tumor tissues isolated from MDA-MB-231 WT cell-injected mice. (**G**) Quantitative analysis of p-IRE1α/t-IRE1α (by Phos-tag), p-JNK/t-JNK, and XBP1s/H3 in (**F**) (n = 6). (**H**) Immunoblotting assay of IRE1α phosphorylation and its downstream signaling pathways in tumor tissues isolated from MDA-MB-231 IRE1α KO cell-injected mice. (**I**) Cell apoptosis in tumor tissues detected by TUNEL staining. Scale bar: 100 μm. (**J**) Quantitative analysis of TUNEL positive area in (**I**) (n = 6). Data are presented as mean ± SEM. Statistical analysis was performed by unpaired Student’s t test. ns, not significant; *p < 0.05; **p < 0.01; ***p < 0.001.

## Discussion

Numerous studies have shown that the UPR, especially IRE1α-XBP1 branch, is activated in tumor tissue and is associated with tumor growth and progression^15,21–24^. Attempts to limit tumor growth by shutting down this pro-survival pathway is quite reasonable and have attracted many researchers, leading to the development of highly selective and potent IRE1α inhibitors^4,25–27^. While attractive and promising, such strategy may have potential shortcomings. First, inhibition of IRE1α-XBP1 cripples adaptive UPR and may aggravate ER stress. This could be detrimental to normal secretory cells that intrinsically experience ER stress and requires IRE1α-XBP1 to maintain the ER homeostasis under physiological conditions. Second, at the dosage that is insufficient to inhibit xbp1 splicing, IRE1α inhibitor may still be able to turn off the pro-death outcome of IRE1α, which usually requires IRE1α hyper-activation and is thus more sensitive to inhibitors. In that case, inhibition of IRE1α may have potential risk in promoting tumor cell growth as it blocks the terminal UPR but keeps adaptive UPR partially functional.

An IRE1α activator may serve as a new strategy to avoid these shortcomings, especially considering that tumor tissue has higher IRE1α activities than normal tissues. Ideally, such activator should not activate IRE1α by itself, but only be effective when cells are already undergoing ER stress and have IRE1α partially activated. Here, we identified Mn^2+^ as a potent IRE1α activator, which luckily meets this goal. Our observation that Mn^2+^ per se does not cause ER stress is different from a previous report in which Mn^2+^ induces ER stress and activates all the three UPR pathways, including IRE1α, possibly due to cell type specificity ^28^. Not only this, Mn^2+^ failed to elevate XBP1 at the protein level, although it promotes *XBP1* splicing. The reason for this still needs further study, and one possibility is that the pro-death outcomes dominates when IRE1α is hyper-activated, leaving no chance for cells to synthesize enough XBP1 protein to survive. Activating the pro-death pathway while leaving the pro-survival outcome of IRE1α unaffected makes Mn^2+^ a promising reagent for cancer treatment. We envisage Mn^2+^-mediated IRE1α activation as an alternative strategy in cancer therapy besides IRE1α inhibition, especially for tumors with intrinsically high level of IRE1α activity.

It remains an interesting question that how Mn^2+^ activates IRE1α. We found that Mn^2+^ interacts with the lumenal domain of IRE1α directly. Like many other kinases, IRE1α requires Mg^2+^ for ATP binding. Mn^2+^ and Mg^2+^ can replace each other in binding with many enzymes, including kinase ^29^. It is possible that Mn^2+^ binds to IRE1α kinase domain similar as Mg^2+^, facilitating ATP binding and phosphate transfer. Notably, Mn^2+^ is more potent than Mg^2+^ in inducing IRE1α auto-phosphorylation (**Figures 3C and 3D**). This may explain why Mn^2+^, but not Mg^2+^, is cytotoxic when cells are experiencing ER stress. Although rare, some kinases do prefer Mn^2+^ over Mg^2+^ as a cofactor, for example casein kinase 2 ^30^. Our study suggests IRE1α to be one of those Mn^2+^-prone kinases. The structural mechanism and physiological relevance of such preference to Mn^2+^ require further study.

Safety of Mn^2+^ usage in cancer therapy should be carefully considered. We did not see cytotoxicity of Mn^2+^ in cell culture. In addition, administration of Mn^2+^ did not cause weight loss in mice. In fact, Mn^2+^-based reagents and chemicals have been widely used in clinic. For example, MnCl_2_ is commonly used for Manganese-enhanced magnetic resonance imaging^31^. Mn^2+^-based nanoparticles have been used in chemodynamic therapy, which accelerate the Fenton reaction to generate hydroxyl radical in tumor microenvironment ^32–34^. Furthermore, Mn^2+^-based nanoparticles could serve as an adjuvant to regulate tumor immune microenvironment and enhance immunotherapy, thanks to the ability of Mn^2+^ in activating cGAS-STING pathway^35^. These examples warrant Mn^2+^ as a safe reagent for cancer therapy. However, one still need to be cautious with long-term Mn^2+^ treatment, as Mn^2+^ can accumulate in the central nervous system and leads to neurotoxicity^36^. Meanwhile, we cannot exclude the possibility that Mn^2+^ induces tumor cell death through binding to proteins other than IRE1α. While our study does not find a specific activator for IRE1α, it does suggest that an ever-been-ignored facet of IRE1α function, in pro-cell death, could be employed and pharmacologically amplified for the purpose of cancer therapy. We look forward to more potent and specific IRE1α activators been developed and practically utilized in the future.

## ACKNOWLEDGMENTS

We are grateful to Prof. Chih-chen Wang and Dr. Lei Wang laboratory (Institute of Biophysics, CAS) who provide MDA-MB-231 cell line. We thank Dr. Yang Yu (Guangzhou Women and Children’s Medical Center; Institute of Biophysics, CAS) for a generous gift of pcW-cas9 plasmid. Thanks to the members of the Wang laboratory for helpful discussions. Acknowledgements are given to Junying Jia (Core Facility, Institute of Biophysics, CAS) for helping with FACS, Ya Wang (Core Facility, Institute of Biophysics, CAS) and Dr. Tao Li (School of Basic Medical Sciences, Peking University) for helping with thermal shift assay, and Chunliu Liu and Yun Feng (Center for Biological Imaging, Institute of Biophysics, CAS) for taking and analyzing confocal images. This work was supported by the National Key R&D Program of China (2021YFA1300800), the National Natural Science Foundation of China (92354303, 32170785), CAS Youth Interdisciplinary Team funding (JCTD-2021-07), and the Strategic Priority Research Program of CAS (XDB37020305).

## AUTHOR CONTRIBUTIONS

Likun Wang designed the study. Ruoxi Shi and Ming Wang performed the majority of the experiments. Si Chen and Xinwei Hu helped with protein purification. Si Chen, Xudong Zhao, Shixin Zhou, Congli Fan, and Zihan Zhou helped with animal experiments. Xudong Zhao and Shixin Zhou helped with sucrose density gradient centrifugation. Ruoxi Shi, Ming Wang, and Likun Wang analyzed the data. Ruoxi Shi and Likun Wang wrote the manuscript.

## STAR Methods

### Key resources table

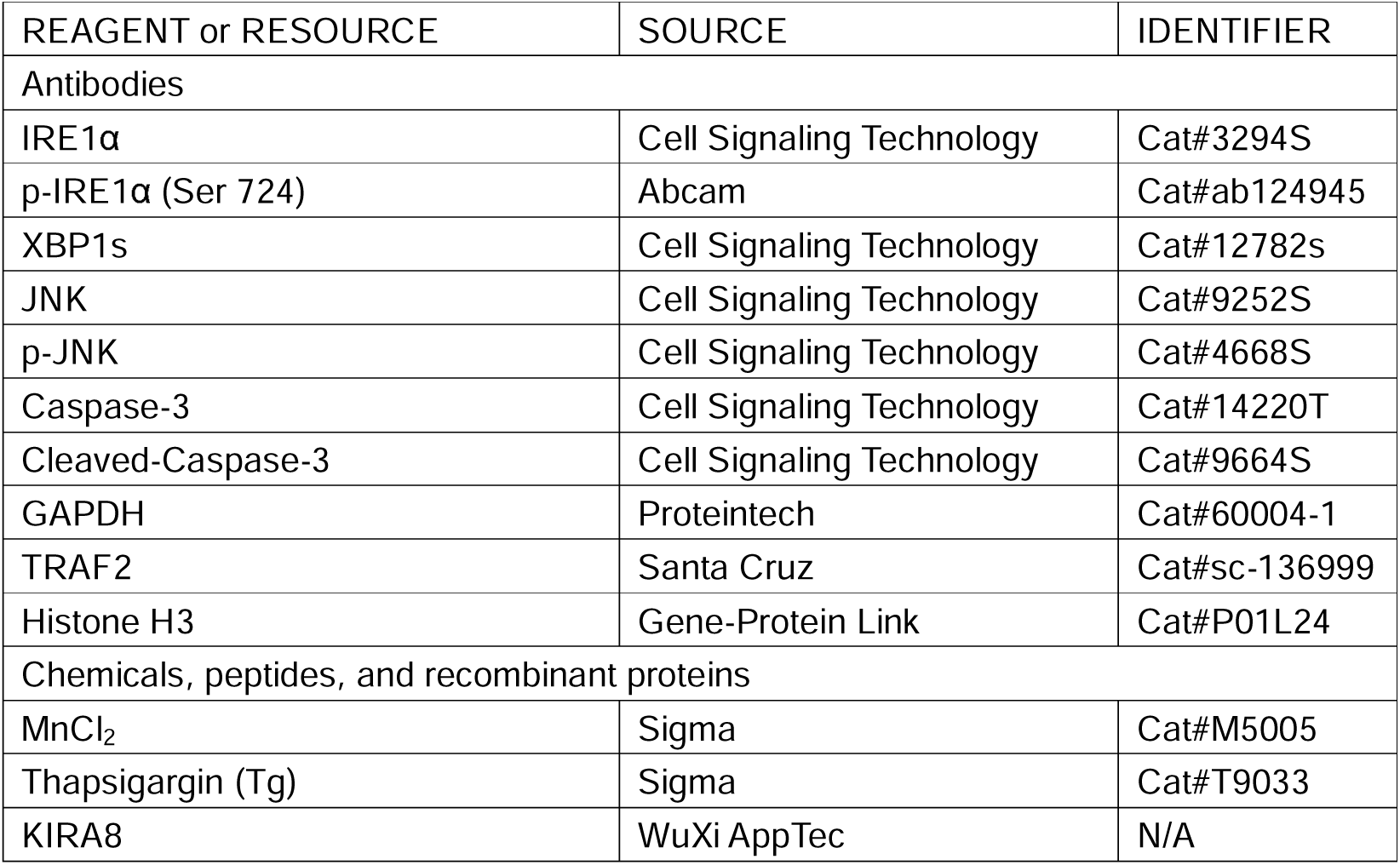
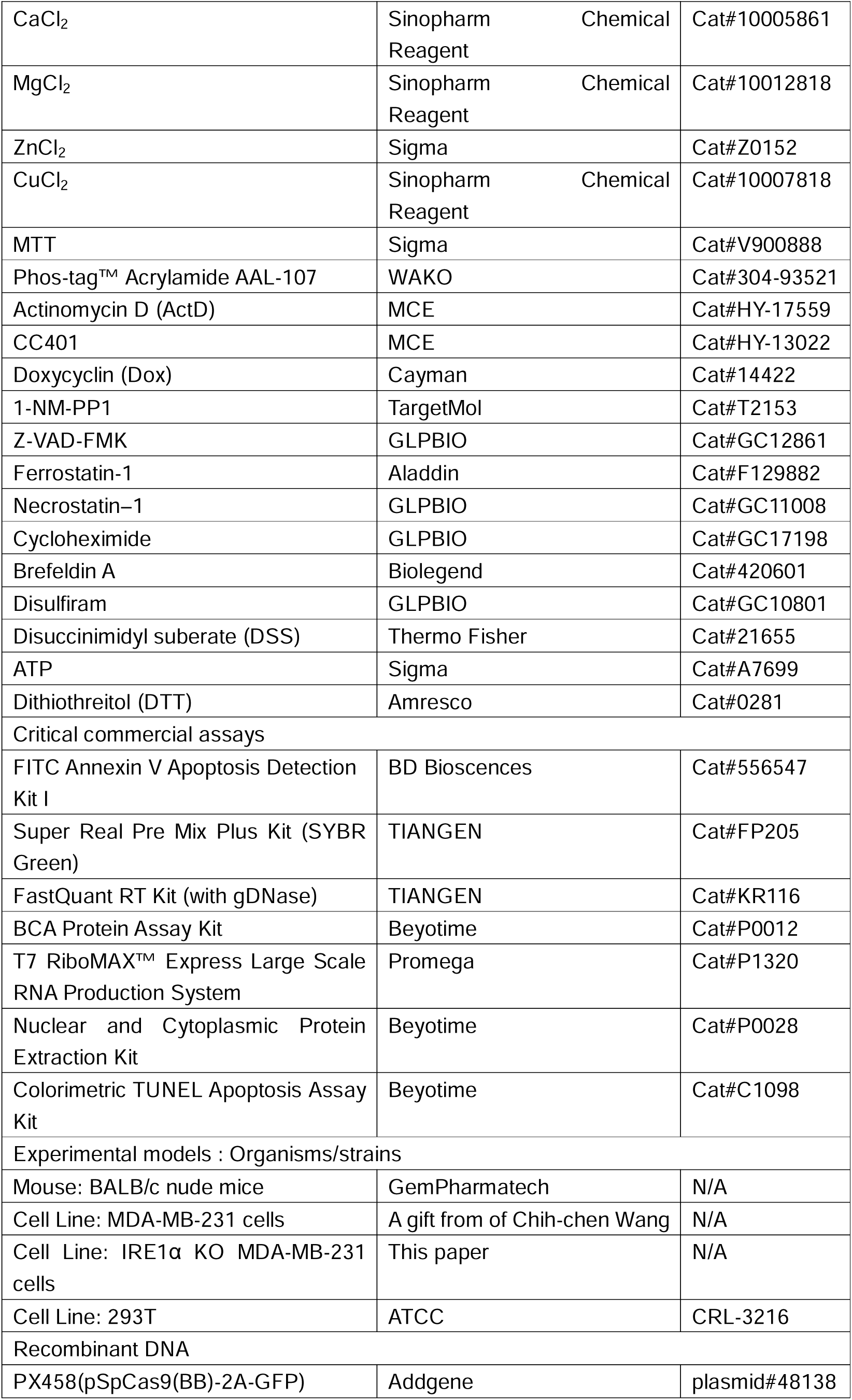
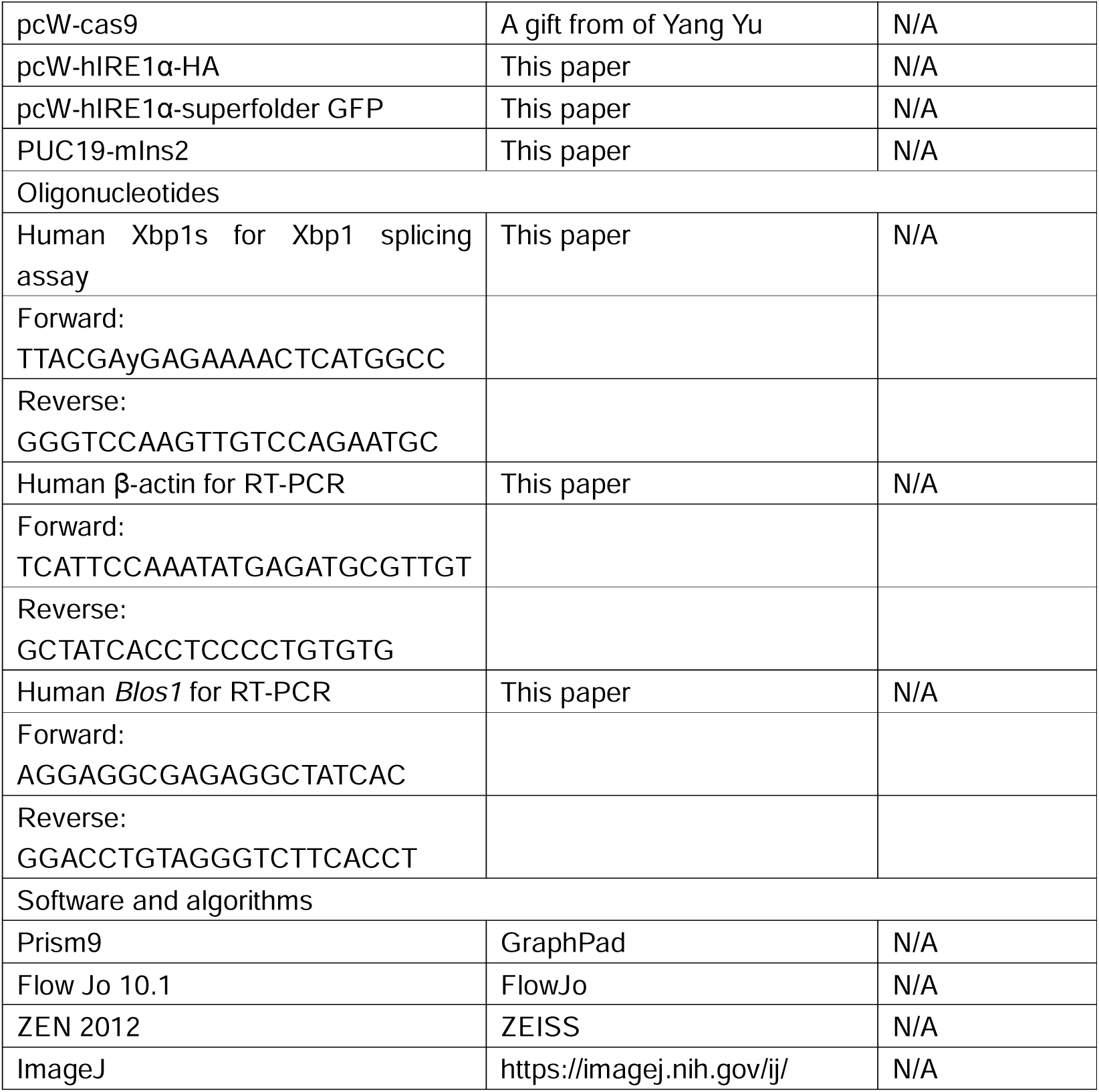

### Animals

BALB/c nude mice were purchased from GemPharmatech. The mice were socially housed (2–5 mice/cage) and given ad libitum access to food and water during a 12-hour light-dark cycle. All animal experiments were approved by the Animal Committee of the Institute of Biophysics, Chinese Academy of Sciences.

For all tumor progression assay, tumor cells suspended in 50 mL of 1 x PBS were mixed with 50 mL of matrigel (BD Biosciences, Cat#356231) and then subcutaneously injected into 6-8 weeks-old mice (1.5 × 10^6^ per mouse). From day 8, peritumoral injections of either saline or MnCllJ (5 mg/kg) were administered every two days. On day 18, mice were euthanized before tumor samples collection. Tumor volume, weight and mice weight were measured and recorded. Tumor sizes were calculated by using the equation V_tumor_ (mm^3^) = a^2^b/2 (a indicates the smallest diameter, b is the perpendicular diameter).

### Cell culture

MDA-MB-231 human breast cancer cells were cultured in Dulbecco’s Modified Eagle Medium (DMEM, Gibco, Cat#C11965500BT) supplemented with 10% fetal bovine serum (Gibco, Cat#10091-148) and 1% penicillin-streptomycin (Solarbio, Cat#p1400). Cells were cultured in an ESCO CO_2_ incubator at 37°C with 5% CO_2_.

For oxygen-glucose deprivation (OGD) assay, cells were washed with PBS, and subsequently, no glucose DMEM (Gibco, Cat#11966025) was added for cell culture. The cells were then cultured in a hypoxic environment with 1% OlJ, 5% COlJ, and 94% NlJ.

For CRISPR/Cas9 knockout cell lines generation, sgRNA targeting human IRE1α(5’-TTGTTTGTGTCAACGCTGGA-3’) was constructed into the vector PX458. Cells were transiently transfected with the appropriate plasmid followed by FACS-based monoclonal selection based on GFP fluorescence. Single knockout clones were verified by immunoblotting and sequencing of the PCR fragments.

The IRE1α reporter cell line and OE IRE1α cell lines were generated by lentiviral infection of MDA-MB-231 IRE1α KO cells with IRE1α reporter (pcW-hIRE1α-superfolder GFP) or IRE1α (pcW-hIRE1α-HA). Stably IRE1α reporter and IRE1α OE cell lines were selected with 10 μg/mL puromycin. Both hIRE1α-superfolder GFP and hIRE1α-HA are expressed in a Dox-inducible way.

### MTT

Cells were plated into a 96-well plate (10³-10^4^ per well). After adhesion, drug treatment was administered. The cells were then incubated at 37°C with 5% COlJ for 24 hours. Following this, 20 μL of MTT (5 mg/mL) solution was added to each well, and incubation was continued at 37°C for 1 - 4 hours. To terminate the incubation, the medium was aspirated and replaced with 150 μL of DMSO (Sigma, Cat#D2650) per well. The plates were agitated at room temperature to dissolve the crystals. Absorbance was measured at 570 nm using a microplate reader, and the results were recorded.

### Flow cytometry assay of cell death

Cell death was measured by FITC Annexin V Apoptosis Detection Kit I (BD Biosciences, Cat#556547) according to the manufacturer’s instruction. Briefly, Aspirate the cell culture medium and transfer it to a 15 mL centrifuge tube. Wash the adherent cells once with PBS and transfer the wash to a centrifuge tube. Add an appropriate amount of trypsin to digest the cells. Incubate at 37°C until gentle tapping can dislodge the adherent cells, at which point digestion should be stopped. Transfer the cells into the centrifuge tube and centrifuge at 1000 rpm for 3 minutes. Wash the cell pellet once with PBS, then centrifuge again at 1000 rpm for 3 minutes. Resuspend the cells in 1× Binding Buffer at a concentration of 1×10^^6^ cells/mL. Transfer 100 μL of the cell suspension (containing 1×10^5^ cells) into a 5 mL flow cytometry tube. Add 5 μL of FITC Annexin V and 5 μL of PI. Gently vortex the cells and incubate at room temperature (25°C) in the dark for 15 minutes. Add 400 μL of 1× Binding Buffer to each tube. Analyze the cells by flow cytometry within 1 hour.

### Immunohistochemistry (IHC) and immunofluorescence (IF)

For live-cell imaging, IRE1α reporter cells were seeded on glass bottom cell culture dishes. Dox (5μg/mL) was used to induce the expression of GFP-labeled IRE1α. Hoechst dye (Beyotime, Cat#C1028) is used for staining cell nuclei.

For tumor tissue TUNEL staining, a Colorimetric TUNEL Apoptosis Assay Kit (Beyotime, Cat#C1098) was used for detect apoptotic cells according to the manufacturer’s protocols. Briefly, paraffin sections were deparaffinized sequentially with xylene, absolute ethanol, 90% ethanol, 70% ethanol, and distilled water. Add 20 μg/mL DNase-free protease K dropwise onto slides and incubate for 30 minutes at room temperature. Wash samples twice with PBS then add 50μl of TUNEL detection solution per sample and incubate 60 minutes at 37℃ in the dark. After mounting samples with Antifade Mounting Medium, observe cells by fluorescence microscopy.

### Immunoblotting assays

For whole cell lysates, remove the culture medium and wash the cells 3 times with PBS. Collect the cells and add RIPA lysis buffer (Beyotime, Cat#P0013B) supplemented with protease inhibitor cocktail (Merck, Cat#539137-10VLCN) and phosphatase inhibitor cocktail (Merck, Cat#524625-1SETCN) on ice for 30 min, then centrifuge at 12,000 rpm for 10 minutes at 4°C. The supernatant contains the total cellular protein.

Nuclear proteins were obtained from cells using the Nuclear and Cytoplasmic Protein Extraction Kit (Beyotime, Cat#P0028) following the manufacturer’s instructions. In brief, scrape cells into a 15mL conical tube and wash them twice with ice-cold PBS. Centrifuge the cells for 3 min at 2000 rpm and discard the supernatant. Resuspend cell pellet in cytoplasmic with ice-cold PBS. Centrifuge the cells for 3 min at 2000 rpm and discard the supernatant. Resuspend cell pellet in cytoplasmic protein extraction buffer A containing protease inhibitor cocktail, vortex vigorously for 5 seconds and then incubate on ice for 10-15 minutes. Centrifuge the lysates at 13,000 rpm for 5 min at 4°C after the addition of cytoplasmic protein extraction buffer B. Carefully remove the cytoplasmic extract from the nuclear pellet. Break nuclear pellet by sonication in ice-cold nuclear extraction buffer containing protease inhibitor cocktail, then incubate on ice for 30 min and centrifuge at 13,000 for 10 min at 4°C. Supernatants were subjected to protein concentration quantification by BCA method and equivalent amounts of proteins were denatured in 5× SDS loading buffer.

Protein concentrations were calculated by the BCA protein assay kit (Beyotime, Cat#P0012). Protein samples in equal amounts were loaded onto SDS-PAGE gel. After electrophoresis, proteins were transferred to a PVDF membrane (Merck, Cat# IPVH00010), blocked with 5% skim milk, and incubated with primary antibodies overnight at 4°C. Then membranes were washed in PBST buffer (136.89 mM NaCl, 2.68 mM KCl, 1.76 mM KH_2_PO_4_, 10mM Na_2_HPO_4_, 0.1% Tween-20, pH 7.2 - 7.4) and incubated with the secondary antibodies at room temperature for 2 h. Finally, the immunoreactive bands were detected with the chemiluminescence reagent (Lablead, Cat#E1050).

Phos-tag SDS-PAGE was performed according to the manufacturer’s protocols. Briefly, 50μM Phos-tag acrylamide (Wako, Cat#304-93521) and 250 μM MnCl_2_ were added into the 5% separating gel when preparing SDS-PAGE. After electrophoresis, the gel was washed with transfer buffer containing 10 mM EDTA (Sinopharm Chemical Reagent, CAT#10009717) for 30 minutes, and proteins were transferred to the PVDF membrane for western blot assay.

### Sucrose density gradient centrifugation

MDA-MB-231 cells were treated with Tg (0.2 μM) and/or MnCl_2_ (100 μM) for 24 hours, then harvested and lysed in lysis buffer. A fresh 20%–40% sucrose gradient was prepared using a gradient maker in 14×89 mm polypropylene tubes (Beckman Coulter). The protein lysates were gently layered onto the gradient. Centrifugation was performed at 35,000 rpm for 20 hours at 4°C using an SW 41 Ti rotor (Beckman Coulter). Approximately 12 mL of solution from each tube was fractionated into 15 fractions from top to bottom. The proteins in the solution were subjected to SDS-PAGE through extraction, and the changes in the IRE1α complex were detected using an IRE1α-specific antibody.

### *Xbp1* mRNA splicing assay and qRT-PCR assays

Total RNA was isolated from whole cells using TRIzol (TIANGEN, Cat#DP424), and equivalent RNA samples were then reverse-transcribed into cDNA using Fast Quant RT Kit (with gDNase) (TIANGEN, Cat#KR116). For XBP1 mRNA splicing assay, cDNAs were PCR amplified with the indicated primers, then resolved on 2.5% (w/v) agarose gels. qRT-PCR employing SuperReal PreMix Plus Kit (SYBR Green) (TIANGEN, Cat#FP205) was performed in the QuantStudio 3 Flex machine (Applied Biosystems) and the QuantStudio 7 Flexmachine (Applied Biosystems), following the manufacturer’s instructions.

### Co-immunoprecipitation (Co-IP)

Cells expressing the target protein were collected and lysed by adding IP lysis buffer (Beyotime, Cat#P0013) on ice for 30 minutes. Subsequently, the lysate was centrifuged at 12,000 rpm for 10 minutes at 4°C. The supernatant was incubated with the primary antibody at 4°C overnight. Following this, Protein A/G Agarose Beads (YEASEN, Cat#36403ES08) were added and incubated for 3 hours at 4°C, after which the beads were washed five times. After adding SDS-PAGE loading buffer, the protein-coated beads were boiled and subjected to SDS-PAGE, followed by detection using Western blotting.

### Lentiviral Infection

HEK293T cells were used for lentivirus packaging. The transfection was performed according to the jetPRIME Transfection Reagent (Poyplus, Cat# PT-114-15) manufacturer’s instructions. Specifically, the viral packaging plasmid psPAX2 (6 μg), pCMV-VSV-G (4 μg), and the target plasmid (10 μg) were mixed in 500 μL of jetPRIME Transfection Buffer, followed by the addition of 40 μL of jetPRIME Transfection Reagent. The mixture was incubated at room temperature for 10 minutes and then added dropwise to the cell culture medium. The virus-containing culture medium was collected 48 hours post-transfection. The medium of the infected cells was removed, and the virus-containing supernatant was mixed with fresh culture medium at a 1:1 ratio for further cell culture. The infection duration was 48 hours, and the fresh virus-containing medium could be replaced during infection. Stable OE IRE1-WT cell lines were selected with 10 μg/mL puromycin in cell culture medium.

### Protein expression and purification

hIREα(CD) (residues 469-977) truncated protein was expressed in SF9 insect cells using the Bac-to-Bac baculovirus expression system (Invitrogen) with an N-terminal 6XHis tag. After 3 days of growth at 27°C, the cells were lysed by sonication in a buffer containing 50 mM HEPES (pH 7.5), 200 mM NaCl, 10 mM imidazole and protease inhibitors. The lysate was centrifuged at 18,000 g for 1 hour at 4°C to remove cell debris and insoluble material. The supernatant was loaded into a pre-equilibrated Ni-affinity column and eluted with lysis buffer containing 250 mM imidazole followed by purification on Superdex 200 column (GE Healthcare). The purified protein was flash-frozen in liquid nitrogen, and stored at 80°C.

The hIREα(LD) (residues 24-441) was expressed in BL21 competent cells using pET-32a plasmid, induced with 0.5 M IPTG at 16°C for 20 hours. After induction, the bacterial culture was centrifuged at 4,000 rpm for 30 minutes at 4°C. The supernatant was discarded, and the cell pellet was resuspended in a buffer containing 25 mM HEPES (pH 7.5) and 500 mM NaCl, supplemented with protease inhibitors. The cells were lysed by sonication, and the lysate was centrifuged at 17,000 g for 1.5 hours at 4°C. The protein was purified using a Ni-affinity column and eluted with buffers containing 10 mM, 20 mM, and 250 mM imidazole. Finally, the protein was concentrated using a 10 kDa cutoff concentrator and further purified by size-exclusion chromatography.

### Thermal shift assay

hIRE1α (CD or LD) and different concentrations of Mncl_2_ were mixed with 10xSYPRO Orange Fluorescent Dye (Sigma, Cat#S5692-500UL) in kinase buffer (20 mM HEPES, 50 mM KCl, 1 mM DTT, 1 mM ATP, pH 7.5). 20 μL mixtures were added into each well of a 96-well qPCR plate (Bio-Rad, Cat#HSP9601) sealed with AB MicroAmp optical adhesive film (Applied Biosystems, Cat#4311971). Protein melting curve was measured by CFX RT-PCR detection systems (Bio-Rad), where the mixtures were incubated at 25°C for 2 min, and then heated in a stepwise fashion from 20°C to 90°C at a rate of 1°C/min.

### In vitro cleavage assay of IRE1α(CD)

Purified hIRE1α(CD), Mn^2+^, and reaction buffer (20 mM HEPES (pH 7.5), 50 mM KCl, 1 mM DTT, and 2 mM ATP) were incubated at room temperature for 20 minutes. Then, 3 μM of mini-XBP1 RNA (5′-FAM-CUGAGUCCGCAGCACUCAG-3′BHQ) was added, and the total reaction volume was adjusted to 30 μL. The reaction was carried out at room temperature for 5 minutes. The reaction was terminated by adding an equal volume of urea, which resulted in a final urea concentration of 4 M. Fluorescence readings were measured using a Microplate Reader (TECAN), with excitation and emission wavelengths of 494 nm and 525 nm, respectively.

### In vitro cross-linking assay of IRE1α (469-977)

10 μM purified hIRE1α(CD), 1 mM ATP, Mn^2+^ (1 or 2 mM), and buffer (20 mM HEPES (pH 7.5), 0.05% Triton X100 (v/v), 50 mM NaCl) were incubated at room temperature for 90 minutes. Then, 250 μM of the cross-linker DSS was added and incubated for 30 minutes at room temperature. The reaction was quenched by adding Tris-HCl (pH 7.5) to a final concentration of 50 mM. The samples were boiled, separated by SDS-PAGE, and immunoblotted with an IRE1α-specific antibody to detect IRE1α(CD).

### In vitro transcription of *mIns2* RNA

The DNA sequence corresponding to *mIns2* RNA (5′-GCCCUCGUCCACUGGAAGUCUGGAACCGUGACCUCCACCGGGUCGUCUUCGCACCGUAACAUCUAGUCACGACGUGGUCGUAGACGAGGGAGAUGGUCGACCUCUUGAU-3′) is inserted into the pUC19 plasmid, and an EcoR1 restriction enzyme recognition site is introduced at the end of the target DNA sequence. The circular pUC19 plasmid containing the target gene is digested with EcoR1 at 37°C overnight. Then the linearized plasmid template is purified using phenol-chloroform extraction. *mIns2* RNA is transcribed at 37°C using the T7 RNA transcription kit. Use DNase I (Roche, Cat#10104159001) to remove the template DNA from the transcription reaction. Finally, the residual DNase I, rNTPs, and other impurities are removed to obtain a purified *mIns2* RNA fragment by ethanol precipitation.

### Data analysis

Statistical analyses were performed using Prism 9.0 (GraphPad). Statistical comparisons between two groups were carried out using unpaired Student’s t-test and the values were shown as mean ± SEM. Throughout all figures: *p < 0.05, **p < 0.01, ***p < 0.001. Significance was concluded at p < 0.05.

---

**Figure S1.**
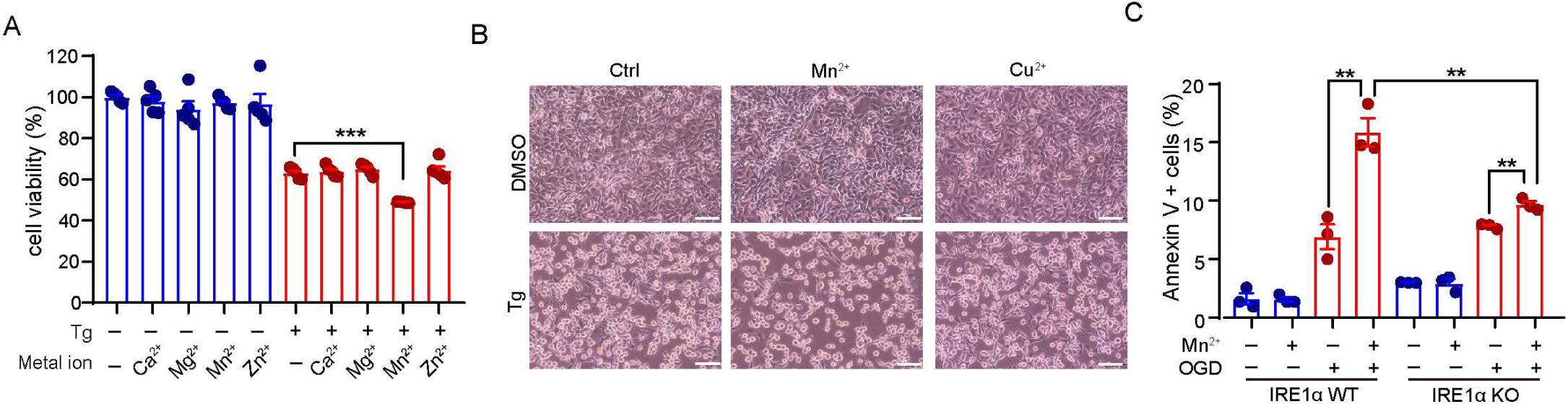
Mn^2+^ aggravates ER stress-induced cell death in an IRE1α-dependent manner, related to Figure 1. (**A**) Cell viability detected by the MTT assay. MDA-MB-231 cells were treated with or without Tg (0.2 μM) in the absence or presence of different metallic compounds (CaCl_2_ (100 μM), MgCl_2_ (100 μM), MnCl_2_ (100 μM) or ZnCl_2_ (100 μM)) for 24 h (n = 5). (**B**) Representative image of MDA-MB-231 cells treated with or without Tg (0.2 μM) in the absence or presence of CuCl_2_ (100 μM) or MnCl_2_ (100 μM) for 24 h. Scale bar: 100 μm. (**C**) Cell death percentage in MDA-MB-231 WT and IRE1α KO cells measured by Annexin V staining. Cells were cultured under physiological conditions or subjected to oxygen-glucose deprivation (OGD) for 8h with or without MnCl_2_ (100μM) (n = 3). Data are presented as mean ± SEM. Statistical analysis was performed by unpaired Student’s t test. ns, not significant; **p < 0.01; ***p < 0.001.

**Figure S2.**
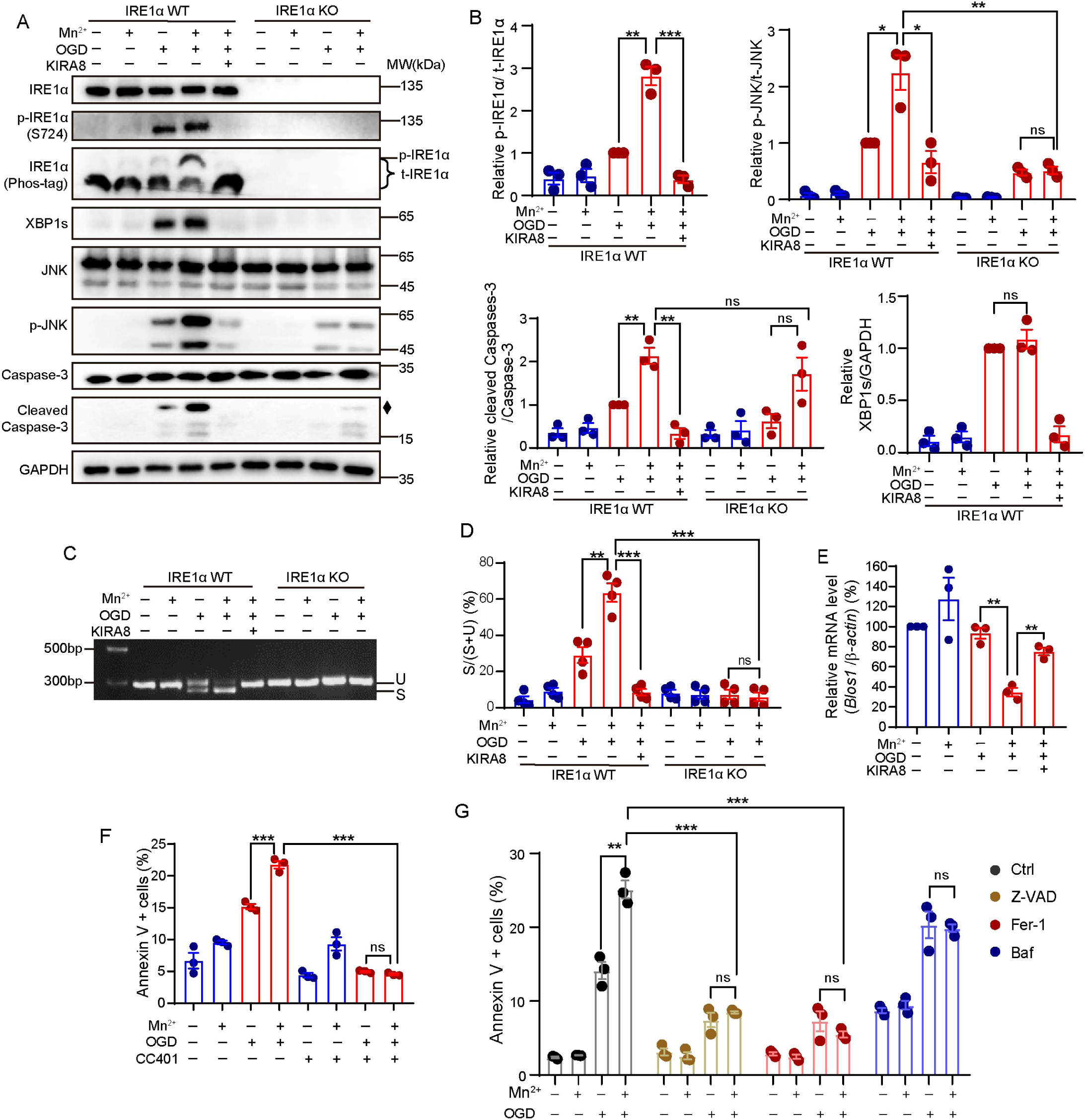
Mn^2+^ promotes the terminal UPR downstream of IRE1α under oxygen-glucose deprivation, related to Figure 2. (**A**) Phosphorylation of IRE1α and activation of its downstream signaling pathway in MDA-MB-231 WT and IRE1α KO cells measured by immunoblotting assay. Cells were cultured under physiological conditions or subjected to oxygen-glucose deprivation (OGD) for 8 h, treated with MnCl_2_ (100 μM) and KIRA8 (1 μM) alone or in combination. Diamond denotes non-specific band. (**B**) Quantitative gray value analysis of p-IRE1α/t-IRE1α (by Phos-tag), p-JNK/t-JNK, XBP1s/GAPDH, and cleaved Caspase-3/Caspase-3 in (**A**) (n = 3). Relative ratio was shown, with ratio in lane 3 (OGD treatment for IRE1α WT) set as 1. (**C**) Agarose gel of *Xbp1* cDNA amplicons from MDA-MB-231 WT and IRE1α KO cells. Cells were treated as in (**A**). U, unspliced *Xbp1*; S, spliced *Xbp1*. (**D**) Quantitative gray value analysis of S/(S+U) in (**C**) (n = 4) (**E**) The mRNA expression level of *Blos1* in MDA-MB-231 cells detected by RT-qPCR. Cells were pretreated with ActD (2 μg/mL) for 1 h, then cells were cultured under physiological conditions or subjected to OGD for 8 h, treated with MnCl_2_ (100 μM) and KIRA8 (1μM) alone or in combination (n = 3). (**F**) Cell death percentage in MDA-MB-231 cells measured by Annexin V staining. Cells were pretreated with or without CC401 (5 μM) for 1 h, then cells were cultured under physiological conditions or subjected to OGD for 8 h (n = 3). (**G**) Cell death percentage in MDA-MB-231 cells measured by Annexin V staining. Cells were pretreated with z-VAD-FMK (z-VAD, 20 μM), ferrostatin-1 (Fer, 1 μM), or bafilomycin A1 (Baf, 10 nM) for 24 h, then cells were cultured under physiological conditions or subjected to OGD for 8 h (n=3). Data are presented as mean ± SEM. Statistical analysis was performed by unpaired Student’s t test. ns, not significant; *p < 0.05; **p < 0.01; ***p < 0.001.

**Figure S3.**
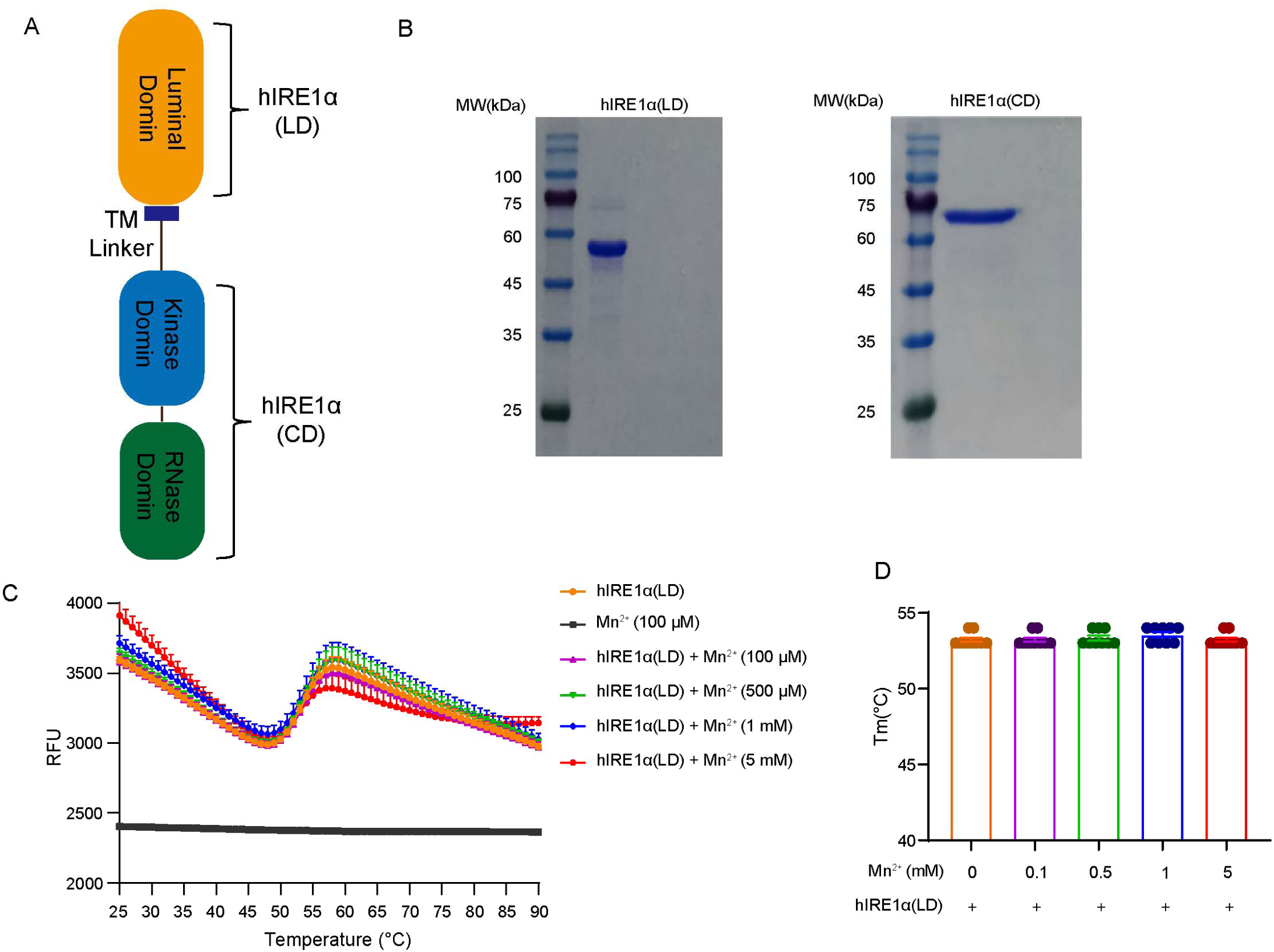
Purification of hIRE1α(CD) and hIRE1α(LD), and the thermal shift assay of hIRE1α(LD) in the presence of MnCl_2_, related to Figure 3. (**A**) Schematic diagram of hIRE1α. (**B**) SDS-PAGE of the purified ER luminal domain (24-441) of hIRE1α (hIRE1α(LD)) and the cytoplasmic domain (469-977) of hIRE1α (hIRE1α(CD)). (**C**) The unfolding curve of recombinant hIRE1α(LD) protein (2μM) incubated with different concentrations of MnCl_2_ measured by thermal shift assay. The melting temperature (Tm) corresponds to the horizontal axis value at the point of maximum slope (n = 9). (**D**) Comparison of the melting temperature of hIRE1α(LD) incubated with MnCl_2_ (n = 9).

